# Cytotoxicity and resistance evolution of a novel antifungal carbon nanoparticle

**DOI:** 10.1101/2024.02.11.579833

**Authors:** Sijan Poudel Sharma, Suraj Paudyal, Justin Domena, Yiqun Zhou, Elliot Cleven, Christian Agatemor, J. David Van Dyken, Roger Leblanc

## Abstract

Antifungal drug resistance is a major problem in healthcare and agriculture. Synthesizing new drugs is one of the major mitigating strategies for overcoming this problem. In this context, carbon-dots (CDs) are a newer category of nanoparticles that have wide applications, potentially including use as antibiotics. However, there is a lack of understanding of the effect of long-term use of CDs as antimicrobials, particularly the ability of microbes to evolve resistance to antibiotic CDs. In this study, we synthesized novel florescent the bottom-up method using two antifungal drugs fluconazole and nourseothricin sulphate (ClonNAT). We first extensively characterized the physical properties of the newly synthesized carbon dots, Flu-Clo CDs. We measured the cytotoxicity of Flu-Clo CDs on budding yeast *Saccharomyces cerevisiae* and determined that it had comparable antifungal inhibition with extensively used drug fluconazole. Furthermore, we demonstrate that Flu-CLO CDs are not cytotoxic to human fibroblasts cell lines. Then, we quantified the ability of yeast to evolve resistance to Flu-Clo CDs. We evolved replicate laboratory yeast populations for 250 generations in the presence of Flu-Clo CDs or aqueous fluconazole. We found that yeast evolved resistance to Flu-Clo CDs and aqueous fluconazole at similar rates. Further, we found that resistance to Flu-Clo CDs conferred cross-resistance to aqueous fluconazole. Overall, the results demonstrate the efficacy of CDs as potential antifungal drugs. We can conclude that yeast populations can adapt quickly to novel antibiotics including CD based antibiotics, including CD-based antibiotics indicating the importance of proper use of antimicrobials in combating infections.

## Introduction

### Fungal infections and resistance

Fungal infections cause more than a billion infections worldwide and 7 million infections in USA (1, 2). Invasive infections caused by pathogenic yeasts have been attributed to mostly *Candida albicans* followed by *Candida glabrata* with *Saccharomyces cerevisiae,* previously considered as non-pathogenic, has also been emerging as opportunistic pathogen (3, 4). With rising cases of fungal infections every year, cases of antifungal drug resistance are emerging as a major healthcare problem leading to high mortality rate among hospitalized patients and increased costs for governments for controlling nosocomial infections. The United States Center for Disease Control (CDC) has stipulated that drug resistant *Candida sps* causes 35,000 infections among hospitalized patients only and has listed *C. auris* as urgent threat and *C. albicans* as serious threat in its 2019 report on antibiotic resistance threat (5).

Fluconazole is one of the most widely used antifungal drugs worldwide for fungal disease treatment, especially candidiasis (6–8) along with aspergillosis and cryptococcal infections (9, 10). It has also been used to treat leishmaniasis (11). Along with clinical use, fluconazole is also used widely in agriculture (12, 13). However, fungal pathogens have been evolving resistance to fluconazole in all settings (14–17). Furthermore, evidence suggests that fungi that evolve resistance to one azole drug are cross-resistant to other azole drugs (18–20). Further, azole-resistant *Aspergillus fumigatus* is also affecting agriculture (21) Overall, drug resistance fungi are not only threatening human health but also food security worldwide. One of the main suggested mitigation strategies for this problem has been discovering new antifungal drugs, as currently there are limited antifungal drugs (22).

### Carbon dots as potential antifungals

Carbon dots (CDs) are an emergent class of nanoparticles with a wide range of applications from drug delivery, cell imaging and ion sensing (23). They are a new type of fluorescent carbon based nanoparticles with sizes ranging from 1∼10 nm (24). They can be synthesized by using various chemicals or precursors including small molecules, polymers, or natural materials (25–27). They are gaining popularity due to their characters such as biocompatibility, non-toxicity, water solubility, photostability (28). New studies have shown that they could be toxic to microbes (29–31), suggesting their potential use as antibiotics. These studies have mostly tested the effectiveness against bacterial species (32). Regarding applications in fungi, many studies have used CDs for imaging purposes (33–35). There are a few studies that have shown some fungal inhibition in *Candida* sps (36–39). The common mechanism of antifungal inhibition in these studies has been doping CDs with toxic metals or other compounds which could potentially be toxic to humans thereby making them unsuitable for medical applications. More research is needed to find safer CD alternatives. In addition, the majority of studies have performed quick cytotoxic assays with the CDs and measure their degree of inhibition. We are unaware of any long-term evolution experiments that assess the ability of microbes to evolve resistance to CDs.

In this study, we synthesized novel antifungal CDs (Flu-Clo CDs) by applying the bottom-up method using a combination of fluconazole, a widely used antifungal drug, and Nourseothricin (ClonNat), a broad range antibiotic as precursors. First, we tested if the newly synthesized Flu-Clo-CDs could inhibit the growth of budding yeast *Saccharomyces cerevisiae*, a widely used model organism. We then tested if Flu -Clo CDs are toxic to human cell lines in order to determine their suitability for medical applications. Next, we tested the ability of yeast to evolve resistance to Flu-Clo CDs using a long-term evolution experiment. We compared these results to those from the widely used antifungal drug fluconazole. Finally, we measured the cost of drug resistance, an important parameter determining the spread of antibiotic resistance in populations and if evolution to Flu-Clo CDs conferred cross resistance to fluconazole. Overall, our study seeks to determine the efficacy of Flu-Clo CDs as antifungals.

## Materials and Methods

### CDs synthesis

We used the bottom-up method for synthesizing CDs. 10mg Fluconazole (Sigma-Aldrich) and 100 mg Nourseothricin sulphate (Research Product International) were dispersed in 15 ml of water to make the molar ratio 1:6 (Fig 1). The mixture was ultrasonicated for 15 min. The solution was transferred to 20 ml autoclave cooled down to room temperature, crude CD solution was obtained. The solution was centrifuged at 3000 rpm for 15 min to remove any precipitates. The solution was transferred to a molecular weight cutoff 100-500 Da dialysis bag and dialyzed with 4 L deionized water for 3 days. Then the solution was concentrated using rotavapor until about 10 ml remained. After that, the solution was subjected to freeze-drying to obtain solid CDs of ≈ 30 mg.

**Fig 1:**
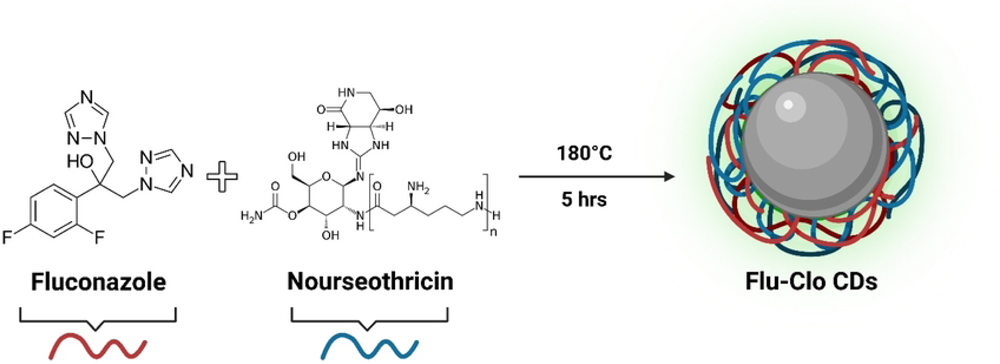
Schematic diagram of synthesis of Flu-Clo CDs. Fluconazole and Nourseothricin sulphate were mixed in 1:6 ratio to synthesize Flu-Clo CDs using bottom up method.

### CDs Characterization

The prepared carbon dots were studied using UV−vis (1 cm cell quartz cuvette) in a Cary 100 UV−vis spectrophotometer (Agilent Technologies) in DI water. The fluorescence spectra of the Flu-Clo CDs were measured in a 1 cm path-length quartz cuvette using a Horiba Jobin Yvon Fluorolog-3 with a slit width of 5 nm for both excitation and emission wavelength. Atomic force microscopy was performed with an Agilent 5420 AFM (Agilent Technologies) using tapping mode, and transmission electron microscopy (TEM) using a JEOL 1200× TEM. Fourier transform infrared (FTIR) spectroscopy was performed using PerkinElner Frontier ATR instrument. Further, we also analyzed Flu-Clo CDs by mass spectrometry. Two techniques (find the technique (names) were used, The mass of the Flu-Clo CDs was estimated to be 635.18 daltons.

### Confocal Microscopy

The objective of performing confocal microscopy was to observe if Flu-Clo CDs can enter *Saccharomyces cerevisiae* cells. For performing the confocal microscopy, we followed the protocol from Seven,et al, 2021. Briefly, overnight saturated culture of yeast was diluted 10-fold with YPD along with Flu-Clo CDs. No antibiotics were dispersed in the control group. The cells were allowed to grow in log phase for 4 hours. After that the culture was washed with PBS and fixed with 4% formaldehyde. Slides were prepared by using 10 ul of cell suspension. The cells were analyzed immediately via confocal microscopy (40).

### Cell viability assay

We performed MTS assay to observe the cytotoxic effects of Flu-Clo CDs in human fibroblast cell line 2091(ATCC) using the procedures described in One Solution Cell Proliferation Assay kit. Briefly, around 5000 cells were seeded in 96 well plates. The following day, the cell culture media was aspirated and new media with treatments in concentration 15mg/lt, 20 mg/lt and 25mg/lt of fluconazole and CDs were added to the adhered cells. Control wells had cells with only media and blank wells had no cells but only media. Next day, 20 ul of the MTS reagent was added to the wells, incubated for 4 hours. Cell viability calculation was performed after measuring the OD at 490 nm. The cell viability was measured by using following formula: (Absorbance of sample/Absorbance of control) *100%

### Cytotoxicity testing of Flu-Clo CDs and fluconazole on *S. cerevisiae*

The experiment was conducted to find if the synthesized CDs inhibit the growth of yeast. Experiment was conducted in 96 well plates. The diploid yeast strain (YPS002) was cultured to saturation following dilution of 1000 folds in 96 well plates with Yeast -Extract Peptone Dextrose (YPD) media. Two concentrations of fluconazole 15 mg/lt and 20mg/lt were used whereas three concentrations of CDs: 15mg/lt, 20 mg/lt and 30 mg/lt were used. One control group with only YPD was used. The plate was incubated at 30 degrees with 6 replicates for each group for 2 days with OD measurements every 15 minutes using Tecan Infinite 200 PRO. Growth rates were calculated using the log phase of the growth curve plotted by using OD readings at 660 nm using previously used custom Matlab code (41, 42).

### Long term evolution experiment

We established replicate *S.cerevisiae* clonal populations in liquid cultures in 96 well plate by inoculating a single colony streaked on a solid agar plate. The plate had 4 treatment groups, two concentrations, 15 and 20 mg/Lt of each fluconazole and Flu-Clo CD with 15 replicates each. Each replicate culture was diluted 2^10 times in fresh media with the antibiotics every 24 hours using Biomek Pro Automation liquid handling robot for 25 days. This protocol results in 10 generations of cell division everyday (43).

For performing competition assays, we froze down day 0 and day 25 plates at −80°C. The plates were thawed and populations from day 0 were competed against day 25 (generation 250) with three sub replicates for each original well population. We found average growth rates for each cell population using the procedure described for cytotoxic assay and performed subsequent analysis for measuring the evolution drug resistance and cross resistance. For statistical analysis, we used JMP Pro 16, Origin Pro 9.1, and Social Science statistics.

## Results

### Flu-Clo CDs synthesis and properties

The synthesized sample of Flu-Clo CDs (3µg.ml^-1^) was tested in a 1 cm optical for UV-vis absorption spectroscopy which showed that maximum absorption at 260 nm and weak band at 295 nm (Fig 2a). The absorption band at 260 nm can be assigned to a π-π* electronic transition of C=C groups and the band at 295 nm can be attributed to n-π* transition of C=O functional groups present in most CDs (44). Furthermore, investigation of photoluminescence (PL) properties shows that the synthesized Flu-Clo CDs show excitation dependent emission while exciting from 250 to 500 nm with emission maxima at 435 nm (Fig 2b). This excitation dependent PL emission can be due to different surface functionalities present on the surface of synthesized CDs which has been explained in previously published articles (45).

**Fig 2:**
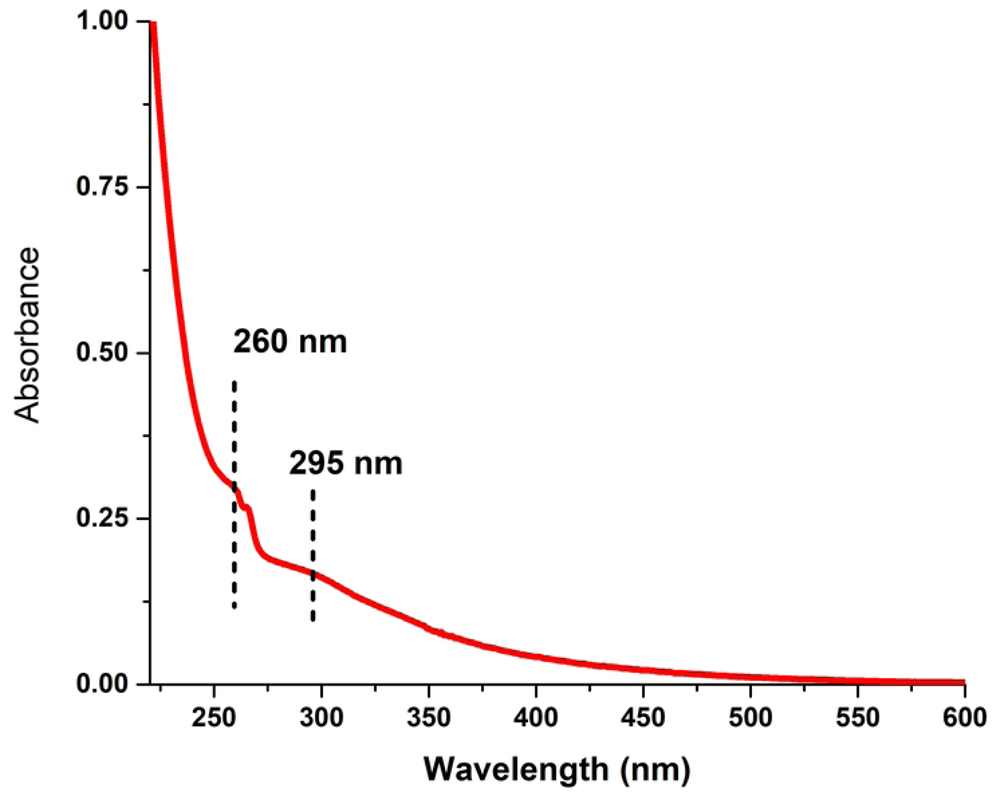
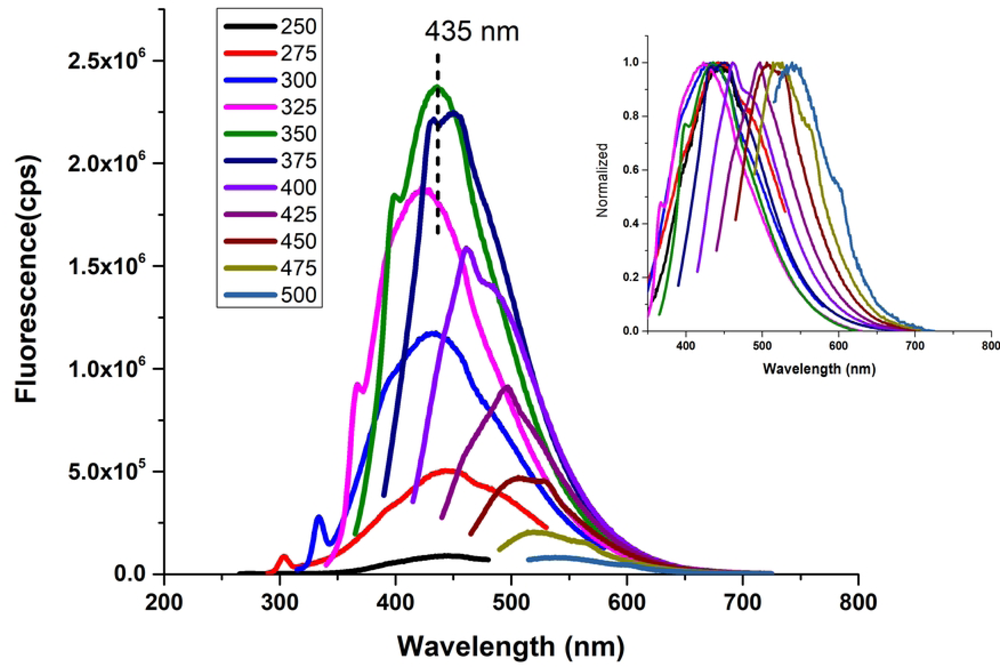
(a) UV-vis absorption spectra of Flu-Clo CDs. Sample (3μg.ml-1) was tested in a 1 cm optical for UV-vis absorption spectroscopy which showed that maximum absorption at 260 nm and weak band at 295 nm. (b) **Normalized photoluminescence emission spectrum of Flu-Clo CDs.** Flu-Clo CDs show excitation dependent emission while exciting from 250 to 500 nm with emission maxima at 435 nm

TEM and AFM analysis of Flu-Clo CDs revealed that their size ranged from 1.5 to 2.5 nm (Figs 3 and 4). The histogram below shows that the size (length and width) of Flu-Clo CDs ranged from 1-5 nm.

**Fig 3:**
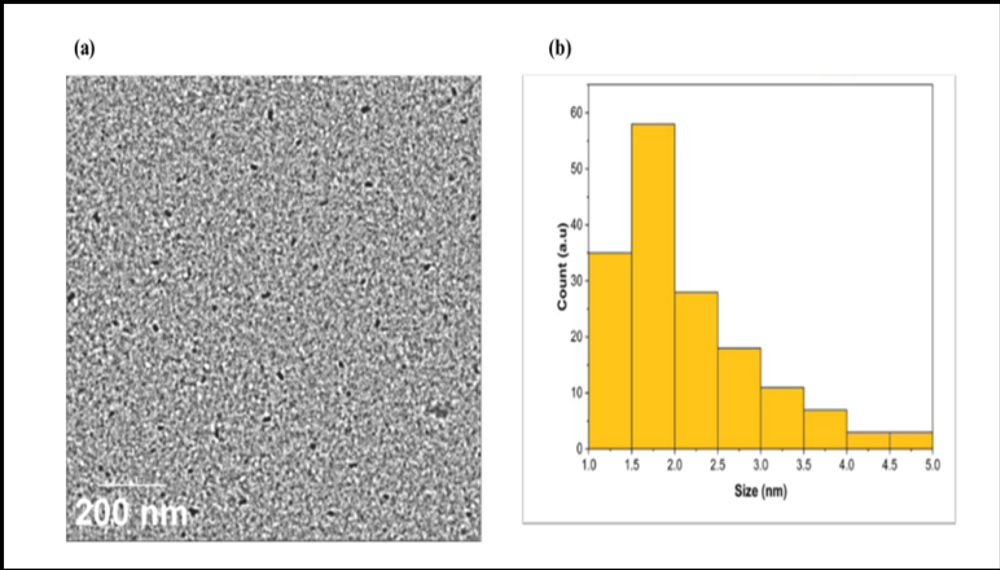
Size analysis of synthesized CDs. (a) Enhanced Flu-Clo CD particles (b) Histogram from TEM showing size range from 1.5 nm to 2.5 nm

**Fig 4:**
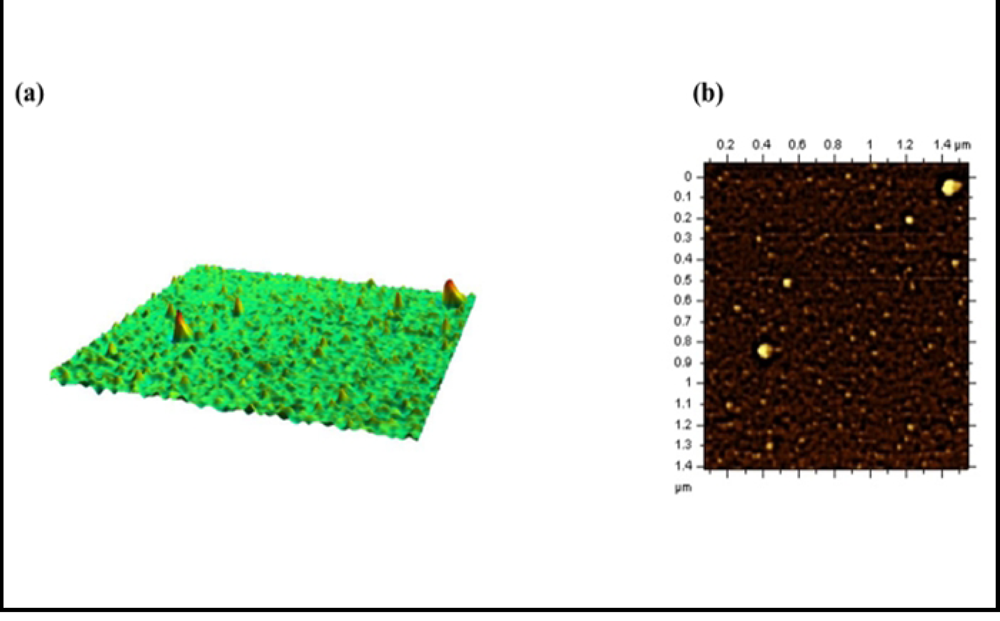
AFM analysis of newly synthesized CDs. (a) 3D image of Flu-Clo CD particles on surface (b) Height determination of Flu-Clo CDs reveal that they are 1 to 5 nm

FTIR-ATR spectroscopy for Flu-Clo CDs was done in solid state to indicate functional groups (Fig 5). It showed a broad vibration band between 3650-3350 cm^-1^ which attributed to O-

**Fig 5:**
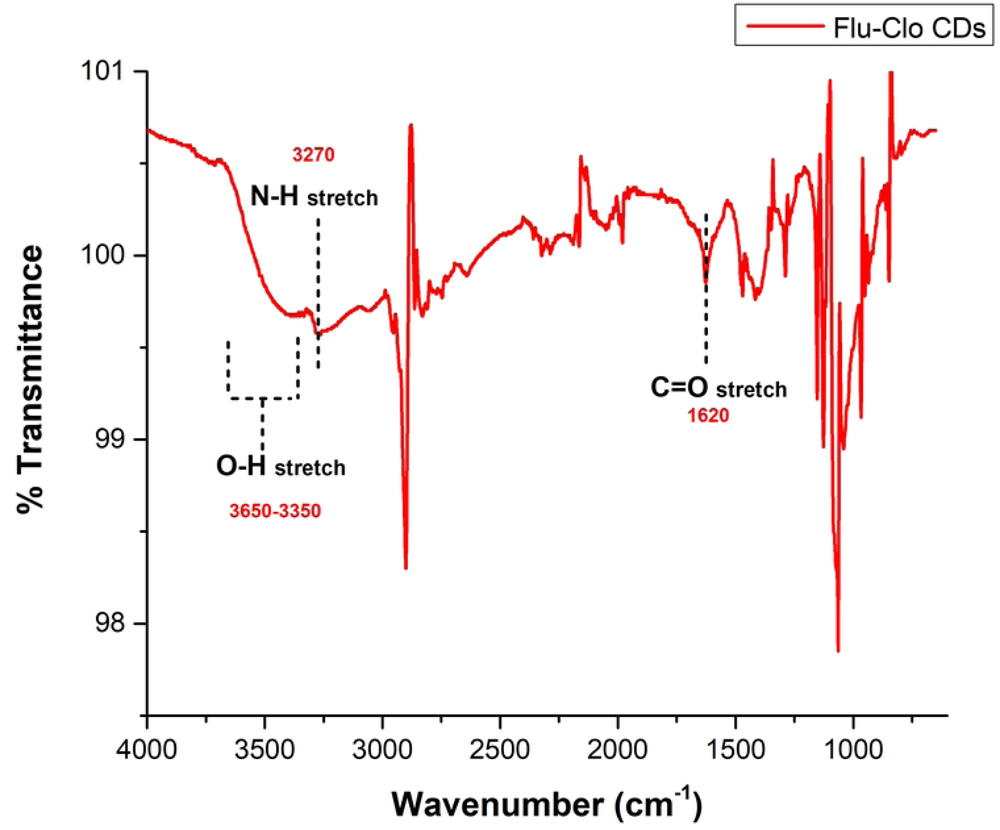
FTIR image of Flu-Clo CDs. FTIR-ATR spectroscopy was performed using PerkinElner Frontier ATR instrument in solid state. Broad vibration bands were observed between 3650-3350 cm-1 (O-H stretch), 3270 cm-1 and 1620 cm-1 indicating O-H stretch, NH stretch, and C-O The C-O stretch at 1620 cm-1 provides evidence of carbonyl functional group in Flu-Clo CDs.

H stretch and a vibration stretch at 3270 cm^-1^ can be attributed can be attributed to N-H stretch. The C-O stretch at 1620 cm^-1^ provides evidence of carbonyl functional group in Flu-Clo CDs. The presence of these functional groups can be explained due to elemental presence of C, N, O in its precursor molecules which are fluconazole and nourseothricin.

### Confocal Microscopy

Confocal microscopy showed that Flu-Clo CDs could enter yeast cells because cells with 15mg/L Flu-Clo CDs showed green florescence but not the cells grown in YPD without Flu-Clo CDs (Fig 6). This shows that the stress caused by toxic CDs to the cells could be because they are able to enter the cells and slow the cell growth. The mechanism by which the CDs were able to enter the cells is not well understood and needs further analysis. However, it could be speculated that it could be like the mechanism of the antibiotics used to synthesize Flu-Clo CDs.

**Fig 6:**
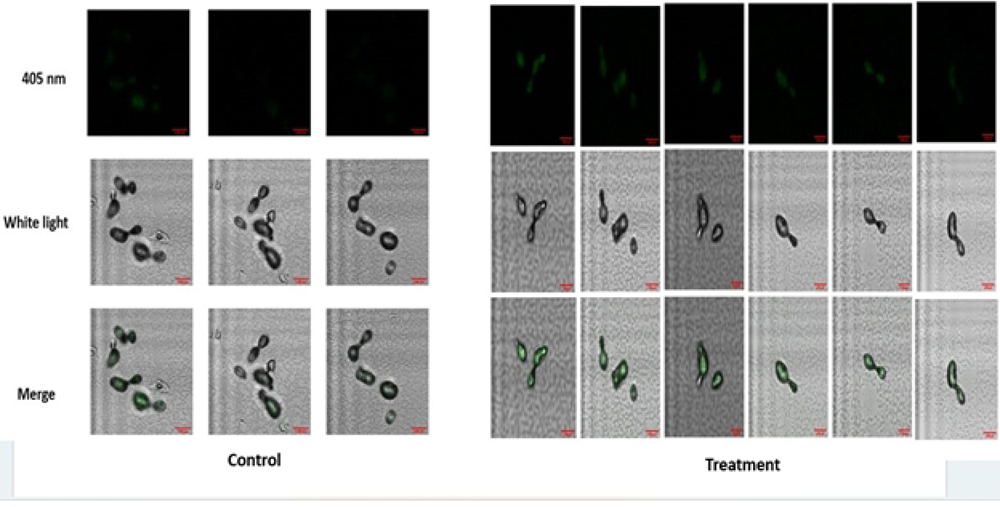
Flu-Clo CDs can enter yeast cells. Confocal microscopy was performed at 405 nm with white light and merge. Left panel: control (no Flu-Clo CDs). Right panel: treatment with 15 mg/L. The objective was 20x while the zoom was 24x. After washing the cells 3 times with PBS to remove autofluorescence, treatment group showed considerable green florescence indicating the presence of Flu-Clo CDs.

### Cytotoxic Assays

We found that the newly synthesized Flu-Clo CDs could remarkably decrease the growth of *S. cerevisiae* (Fig 7). The results show that 15 and 20mg/lt aqueous fluconazole inhibited mean growth rate by 17.5% and 27.5% respectively compared to YPD. Flu-Clo CDs 15, 20 and 30 mg/L reduced the mean growth rate by 26.8%, 28.9 %, 21.7%and 24.06% respectively. We observed that fluconazole has high variability on yeast inhibition across the replicates consistent with previous experiments. One way resulted in significant difference in growth rates across treatments (F=3.7, p value 0.02). Pairwise Tukey HSD test revealed that that compared to control (YPD), fluconazole 20 mg/L and Flu-Clo CDs 15 and 20 and 30mg/L were able to reduce the growth rate significantly (p<0.05) whereas fluconazole 15mg/Lt inhibition was not statistically significant. However, this method was not compatible to compare the toxicity with ClonNat in same concentrations in YPD as these concentrations of drugs were lethal to *S. cerevisiae*, not semi lethal as the concentrations of other two drugs used in this experiment.

**Fig 7:**
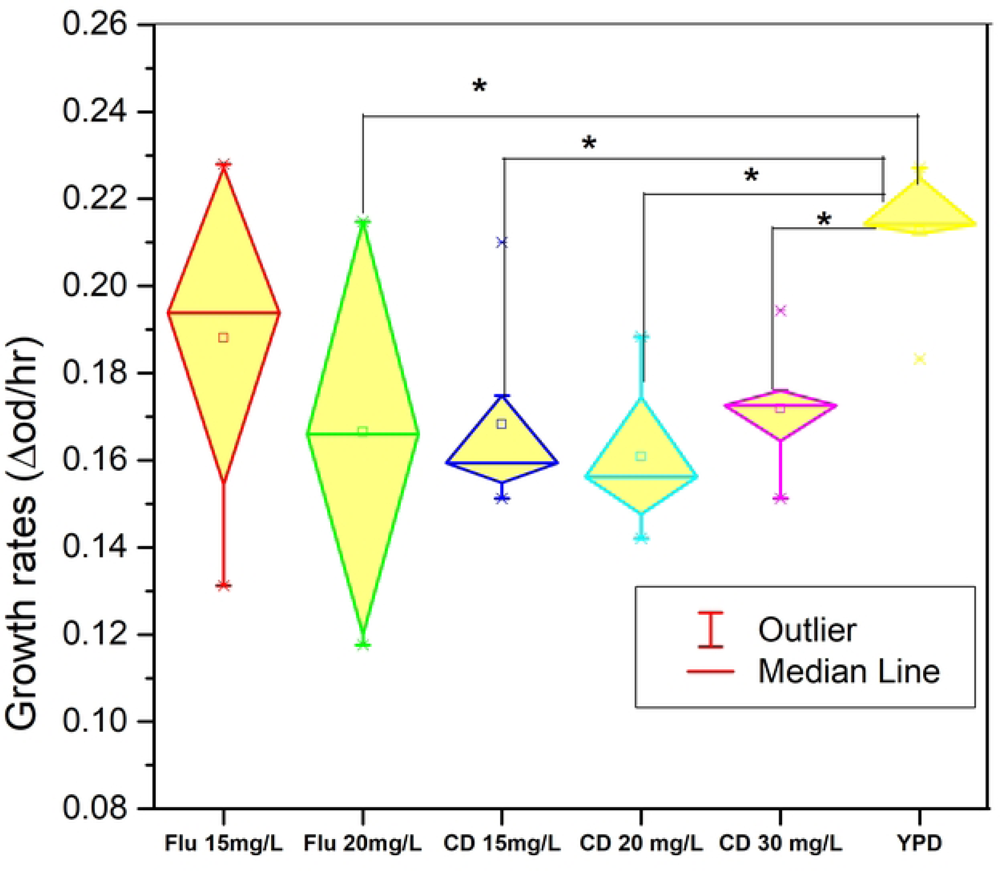
Flu-Clo CDs can inhibit growth of budding yeast *S. cerevisiae*. Growth rates (Rmax) of yeast in YPD with fluconazole 15 and 20mg/L and Flu-Clo CDs 15, 20, 30 mg/L. Growth rates were calculated using the log phase of the growth curve plotted by using OD readings at 660 nm using previously used custom Matlab code. One way ANOVA test showed significant difference among groups (p<0.05).

We wanted to understand if the newly synthesized Flu-Clo CDs were toxic to human cell lines. We did so by performing a cell viability assay MTS assay that measures the degree of cell proliferation. We used 15, 20 and 25 mg/L of both fluconazole and Flu-Clo CDs (Fig 8). After taking readings and performing One-way ANOVA, we found that there was no significant difference among the groups(F>0.17847, p>0.05). Tukey’s pairwise test showed no significant difference between any experimental groups with the median cell viability above 100%. for all groups. Therefore, both drug groups in the concentrations used were not toxic to human fibroblast cell lines.

**Fig 8.**
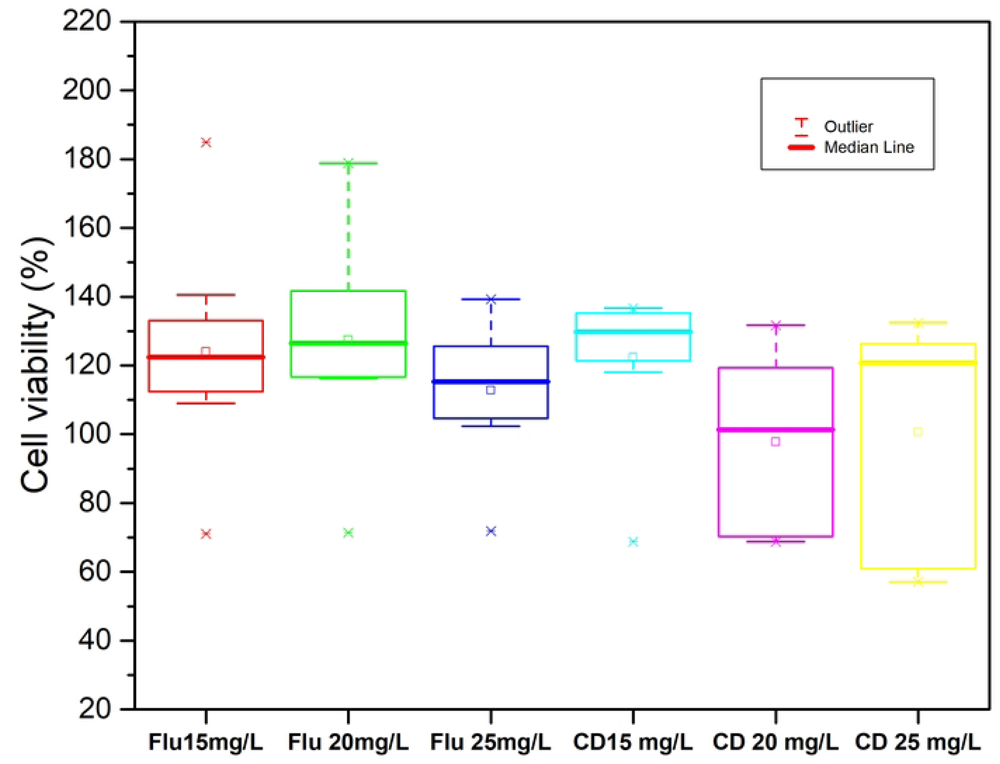
In vitro 2091 Cell viability percentage measured by MTS assay. Three concentrations, 15, 20 and 25mg/L was used for both the drugs. All groups had considerably high cell viability with median values more than 100%. One way ANOVA showed no difference in the cell viability among groups and pairwise comparison using Tukey Test also showed no difference in viability.

### Long term evolution with Flu-Clo CDs and fluconazole

To compare how Flu-Clo CDs and fluconazole differ in their susceptibility to evolutionary resistance, we employed experimental microbial evolution using *S. cerevisiae* exposed to different concentrations of these drugs. We measured the growth rates of beginning and end point populations after 250 generations of experimental evolution in presence of drugs and calculated the percentage difference to obtain the change in population fitness over time. We found that the average fitness of all the four groups’ end point populations increased over time (Fig 9). For fluconazole 15 and 20 mg/L (FLU15 and FLU20), the average increase in fitness were approximately 31% and 36% respectively and for Flu-Clo CD 15 and 20 mg/L (CD15 and CD20), they were approx.49% and 53% respectively. One way ANOVA revealed that there was no significant difference across the groups (P=0.306) for percentage change in fitness after 250 generations of experimental evolution. We further performed Tukey HSD test for pairwise comparison which also showed no difference at 0.5 level between the means of any treatment pairs. High variance was observed in all groups but were not significantly different at 0.05 alpha level. Most replicates in each group had increased fitness whereas few populations had decreased fitness over time in each group.

**Fig 9:**
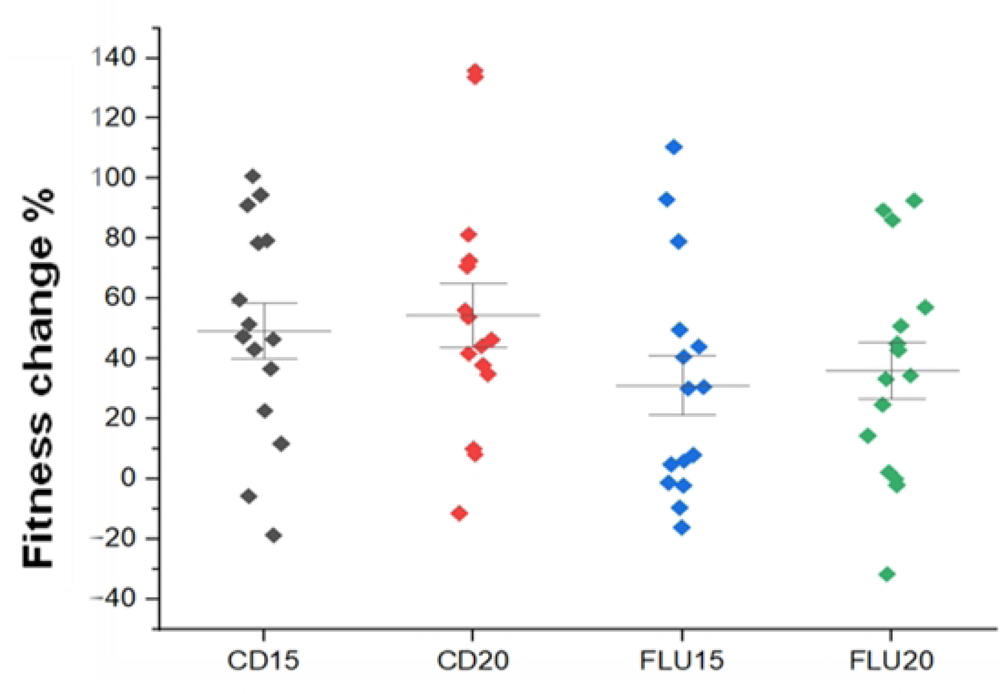
Evolution of resistance of *S.cerevisiae* population to Flu-Clo CDs and fluconazole after 250 generations. Experiment was conducted in 96 well plates with four treatment groups, two concentrations, 15 and 20 mg/L of each fluconazole and Flu-Clo CD. Evolution of resistance was measured by calculating percentage change in fitness (growth rate) over 250 generations. One way ANOVA revealed that there was no significant difference across the groups (P=0.306) (n=15)

**Fig 10.**
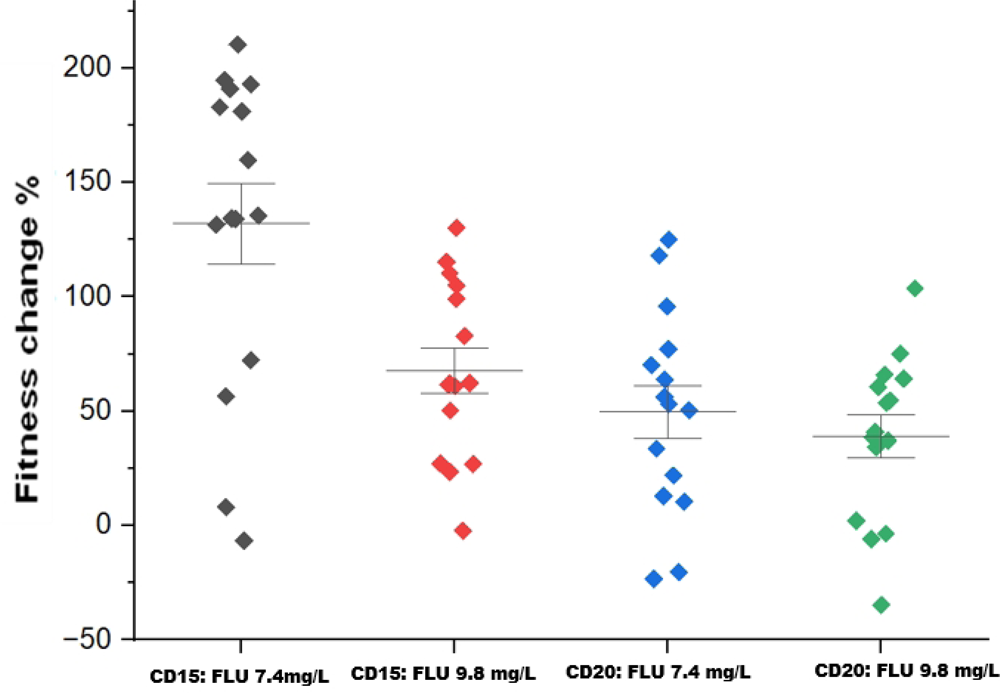
Evolution of cross resistance in evolved lines to fluconazole. Percentage change in fitness of populations evolved in Flu-Clo CDs in same and different molarities of fluconazole. Flu-Clo CDs in concentration 15g/L labelled as CD15 in figure was estimated same molarity as fluconazole 7.4mg/L (0.000024M) and Flu-Clo CD 20 mg/L labelled as CD20 in the figure had same molarity as 9.8 mg/L (0.000032M). The beginning and evolved yeast population in Flu-Clo CDs were thawed and diluted in fresh media with fluconazole and the difference in growth rates were measured.

Next, we investigated if evolution if evolution of resistance to Flu-Clo CDs used conferred cross-resistance to fluconazole. We observed that growth rates of most populations in 250^th^ generations were remarkably higher than that of the beginning population (Fig10). We calculated the percentage increase or decrease in fitness of populations that evolved on 0.000024M and 0.000032M Flu-Clo-CDs in the same molarities of fluconazole. Yeast populations adapted to Flu-Clo CDs were able to adapt to fluconazole in the same molarity and different molarities. Like our previous experiment, divergence in population fitness was observed in wells with fluconazole.

For understand if any cost had occurred in the end point populations resulting from the adaptation, we also investigated if there was any fitness change in YPD a nutrient rich media without the antibiotics. We observed a range of changes in fitness (Fig 11). Most end point populations had increased fitness in YPD compared to beginning population as well indicating no fitness cost to adaptation to antifungals similar to previous studies conducted in *S. cerevisiae* (46). One way ANOVA followed by Tukey test for pairwise comparison revealed no significant difference in fitness change among the treatments (p>0.05, n=15). While all replicate populations from CD15 had an increase in fitness, 33.33% of FLU15 group, 40% of FLU20 and 26.67% of CD20 group had a decrease in fitness in YPD compared to the starting population. This indicates that there could be tradeoffs to adaptation to drugs causing reduced fitness in previously well adapted environment.

**Fig 11.**
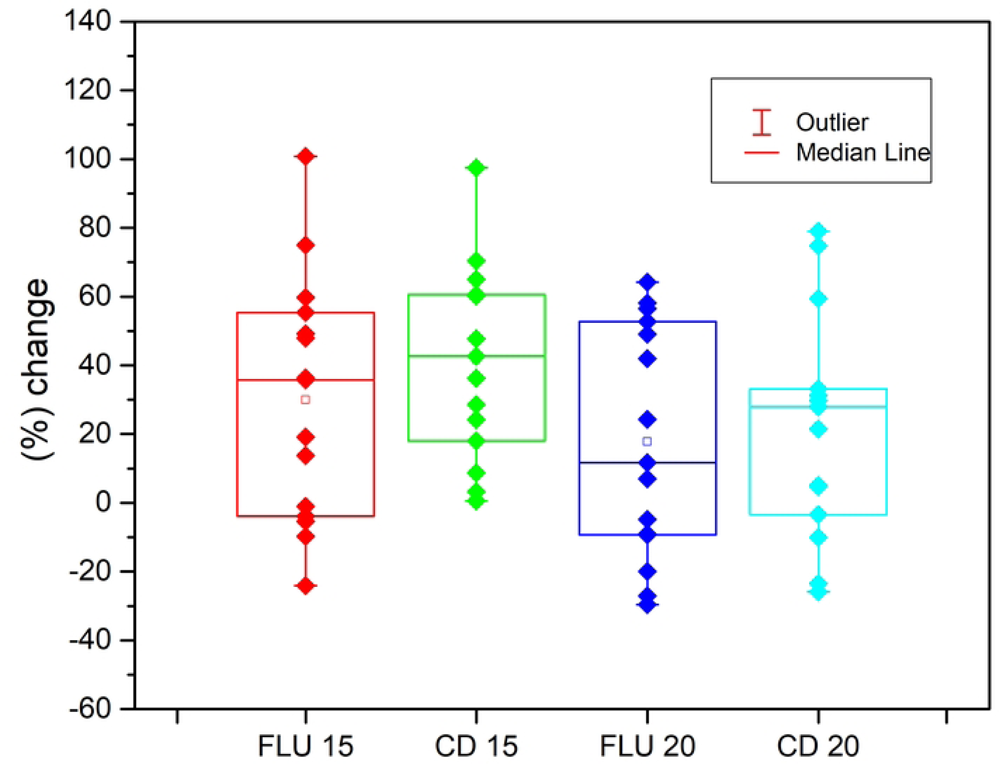
Percentage change in fitness in nutrient-rich environment (YPD) after 250 generations. After evolving in fluconazole 15mg/L and 20 mg/L (FLU 15 and FLU 20) and Flu-Clo CDs 15mg/L and 20 mg/L respectively, their growth rates were measured in YPD and percentage difference in fitness was analyzed to find if there was any cost of resistance. Majority populations did not have any trade off from evolving in antibiotics. One way ANOVA showed no difference between the treatment groups (p>0.05, n=15)..

## Discussion and conclusion

Azoles, polyenes and echinocandins are three classes fungal drugs that are traditionally used in clinical medicine and agriculture. However, the problem of emergence of worldwide antifungal drug resistance is rising, threatening public health and agriculture (22). One of the ways suggested to mitigate the problem is the development of new drugs. Recent studies have shown that Flu-Clo CDs can be effective alternative to traditional drugs and can used in multiple ways as anti-microbials such as in combination with other drugs (47) or doping with chemical agents on the surface (48). However, there is a lack of CD based antifungals which are safer to use for medical purposes.

In this study we have synthesized new CD based antifungal drug Flu-Clo CDs using bottom-up method which has been more popular for synthesizing anti-microbial CDs in recent years. (31). This process is advantageous as it is a speedy process, is less costly and is environment friendly (49) although it can cause low yield of CDs. Very few studies have used existing antibiotics to synthesize CDs using drugs such as levofloxacin, ampicillin, kanamycin and metronidazole (50). The synthesized CDs have been mostly targeted against bacterial species. Studies targeting fungal pathogens have used other techniques besides using the antifungal agents themselves. One of the techniques is doping with heteroatoms such as boron, sulfur, nitrogen, iodine, etc (36, 38). Another method includes conjugating CDs with metals such as gold (39) and using both toxic and non-toxic chemicals as precursors for synthesizing antifungal CDs (51–53). However, most tests have used high concentration of drugs and have only performed short term cytotoxic assays.

The newly synthesized Flu-Clo CDs had ideal size of 1 to 5 nm as demonstrated by AFM and TEM making them permeable for entering live cells. They were florescent (Fig 2) and gave green fluorescence to *S. cerevisiae* cells (Fig 6). By performing confocal microscopy, we were able to confirm that Flu-Clo CDs can enter yeast cells, hence it can be speculated that Flu-Clo CDs could act by targeting both cell wall or cell membrane or intracellular organelles although further research needs to be done to confirm each of the hypothesis. This is in line with some other studies where CDs work by photodynamic inactivation (54, 55). Further, because their fluorescent property they can also be used for bioimaging of fungal cells and can be tested to other microbial and animal models similar to previous studies (56, 57).

Regarding the cytotoxic effects of Flu-Clo CDs, we tested the cytotoxic effects in different mass/volume concentrations by measuring the logarithmic growth rates and comparing with fluconazole traditionally used to treat yeast infections and is also a precursor. The results showed that the synthesized CDs were more effective in 15 and 20 mg/Lt and had similar cytotoxicity with fluconazole in same mass/ volume concentrations and higher concentrations did not confer higher inhibition, Most studies use minimum inhibitory concentration (MIC) approach to measure the toxicity of CDs on microbes. (58, 59). A study conducted by Priyadarshini, Rawat et al. 2018 on *C. albicans,* growth curve analysis was performed in different concentrations like our study (39). The results, however, were different as higher concentration of CDs resulted in higher inhibition whereas in our study higher concentration of Flu-Clo CDs did not necessarily cause higher inhibition of growth rate. The concentration of CDs to be used for inhibition may depend on the type of caron dots synthesized and microbe targeted. Another study that that have found good efficacy of CDs in lower concentrations were tested in viruses (60). Further, Flu-Clo CDs maintained their toxicity for months after synthesis showing that they were highly stable.

Another advantage of these Flu-Clo CDs were that these CDs were not toxic to mammalian cells (Fig 8) rendering them safe alternative as potential (61) therapeutic agents, an important aspect often overlooked in CD based studies. This is very important because many potential antifungals may have homologs of similar function in the host, resulting in the possibility of severe side effects in the hosts. Future analysis for this study includes determining the MICs of Flu-Clo CDs. Further work is also necessary to understand the exact mechanism of inhibition for yeast cells. Future steps also include testing the cytotoxicity Flu-Clo CDs in other yeast species like *Candida* using a similar approach.

The results from long term evolution experiment (Fig 9) showed that replicate populations were able to evolve resistance to the antifungals after 250 generations of growing under the antifungal stress. Although, we did not observe any considerable cost of resistance. This could be either because the resistance mutations are cost-free (62, 63) or because 250 generations was enough time for populations to evolve compensatory mutations to eliminate cost of resistance. This finding is much the same as most laboratory studies where increase in fitness is observed in response to prolonged growth under antibiotic stress (64, 65). The mode of adaptation to both drugs can be attributed to local evolution in each well population and not horizontal transfer of resistance gene, a common phenomenon that occurs in bacteria but not in fungi (65). While fitness increased in most replicate populations, there was a range of fitness change with some populations having lower fitness than the original populations indicating that not all populations were able to adapt to the stressed environment. This result is similar to a study conducted by Cowen et al, 2014 where *C. albicans* populations evolved under fluconazole stress diverged in fitness owing to the multiple mechanism of drug resistance (66). Some possible explanation for decrease in fitness can be explained by selective sweeps of deleterious mutations and maladaptive epistatic interactions of mutations (67). However, exact mechanism of resistance or tolerance to carbon nanoparticles have yet to be explored because of the lack of similar studies where long-term studies for effectiveness are conducted. Further, our results indicated that yeast populations adapted to Flu-Clo CDs have potential to develop cross resistance to other drugs. This is not a unique observation as many studies conducted in the past with antibiotics including other nanoparticles have seen similar results in other microbial systems as well. (Chuanchen, Beinlich et al. 2001, Sommer, Munck et al. 2017 (68).

In this way, we have utilized long term evolution experiment to understand the evolution of drug tolerance to both routinely used antifungal fluconazole and novel antifungals Flu-Clo CDs. We recommend this approach for expanding the knowledge on effectiveness of new drugs before dissemination in the market because the molecular mechanisms of resistance arising in experimental populations of both pathogenic and nonpathogenic yeast populations have been found to be similar in natural populations (46).We conclude that new category of drugs, carbon-based quantum dots or CDs could also prone to resistance as a result of continued use. Future work could also include studying the levels of drug resistance when combination of new drug is used with prevalent drugs as this was not tested in this study.

## Acknowledgements

We would like to thank the Department of Biology and Department of Chemistry for providing us with the equipment and expertise for carrying out the experiments.

